# Inferior parietal lobule and early visual areas support elicitation of individualized meanings during narrative listening

**DOI:** 10.1101/301812

**Authors:** S. Saalasti, J. Alho, M. Bar, E. Glerean, T. Honkela, M. Kauppila, M. Sams, I. P. Jääskeläinen

## Abstract

When listening to a narrative, the verbal expressions translate into meanings and flow of mental imagery, at best vividly immersing the keen listener into the sights, sounds, scents, objects, actions, and events in the story. However, the same narrative can be heard quite differently based on differences in listeners’ previous experiences and knowledge, as the semantics and mental imagery elicited by words and phrases in the story vary extensively between any given two individuals. Here, we capitalized on such inter-individual differences to disclose brain regions that support transformation of narrative into individualized propositional meanings and associated mental imagery by analyzing brain activity associated with behaviorally-assessed individual meanings elicited by a narrative. Sixteen subjects listed words best describing what had come to their minds during each 3–5 sec segment of an eight-minute narrative that they listened during fMRI of brain hemodynamic activity. Similarities in these word listings between subjects, estimated using latent-semantic analysis combined with WordNet knowledge, predicted similarities in brain hemodynamic activity in supramarginal and angular gyri as well as in cuneus. Our results demonstrate how inter-individual differences in semantic representations can be measured and utilized to identify specific brain regions that support the elicitation of individual propositional meanings and the associated mental imagery when one listens to a narrative.

## Introduction

When listening to a narrative, the verbal expressions translate into propositional meanings (*i.e.*, semantics) along with the associated mental imagery, with the keen listener seeing with his/her “mind’s eye” the objects, environments, actions, and events in the story. The intriguing question of how the human brain codes the semantics of language has been under investigation for decades. Brain areas sensitive to word meanings have been observed in temporal, parietal and frontal cortices^1,2^. It has been suggested that inferior parietal regions act as convergence zones for concept and event knowledge, receiving input from sensory, action, and emotion systems^1^. Recently, in a study where word-meaning categories occurring in a narrative were mapped onto human cerebral cortex using functional magnetic resonance imaging (fMRI)^3^, the results both agreed with previous meta-analysis of semantic areas of the human brain^1^ and extended our understanding by disclosing both regional selectivity, and the extent of activation responding, to semantic categories. The semantic representations were not confined to left hemisphere, but were observed predominantly bilaterally^3^. However, information is represented in human brain in multiple ways^4^, and listening to a captivating story may, in addition to linguistic semantics, also activate processes related to mental imagery as one sees events with the “mind’s eye”.^5,6^ Previous empirical evidence suggests that when a person forms mental imagery visual cortical areas are activated, which are also the first cortical areas to receive real visual signal from the eyes^4,7^, though there are differences between individuals in the strength of visual imagery^8^.

What previous studies have not yet addressed is that stories can be experienced quite differently^9^ based on differences in previous experiences^10^, e.g., upon hearing the word “dog” one person can come to think of a happy Collie, another an angry Rottweiler. Given such inter-individual differences, we hypothesized that by analyzing brain activity based on behaviorally assessed individual semantics^11^ elicited by a narrative we can disclose brain regions supporting the elicitation of individual semantics and mental imagery during story listening. We presented an eight-minute narrative describing daily events in a woman’s life to 16 healthy females during 3T-fMRI, and asked subjects to report, by listing descriptive words, what had come to their minds while listening to the narrative. We then utilized latent semantic analysis^12^ (LSA) WordNet^13,14^ to quantify similarities-differences in these word listings between each pair of subjects and tested, using representational similarity analysis^15^ (RSA), whether similarities/differences in the individualized meanings predicted similarities-differences in brain activity as quantified using inter-subject correlations^16,17^. We specifically hypothesized to see involvement of brain areas such as inferior parietal lobe and visual cortical areas identified in previous studies as core semantic processing areas^1^ and areas activated during mental imagery^4^. Further, by demonstrating how inter-individual differences in semantic representations can be measured and utilized to map the semantic areas in the brain, our findings also provide an important methodological extension for studying the human semantic system.

## Results

Behavioral responses of the subjects revealed that while some individuals perceived the story semantically highly similarly, there were also robust differences between many subjects in how they heard the story as disclosed by LSA^12^ combined with WordNet^13,14^ knowledge^18^ (**Fig. 1**). Inter-subject correlation (ISC)^16,17^ of brain activity during listening the narrative (**Fig. 2**) was statistically significant (FDR corrected q < 0.05) in an extensive set of brain areas: bilateral frontal (superior, middle and inferior frontal gyri), temporo-parietal (superior, middle and inferior temporal gyri) brain areas, extending also to midline regions such as precuneus and cuneus and right cerebellum. RSA^15^ showed that between-subject similarities in perceived semantics of the story predicted between-subject similarities in local brain hemodynamic activity. Subject pairs whose individual semantics were similar also exhibited similar brain activity in bilateral supramarginal and angular gyrus of the inferior parietal lobe, and in the occipital pole (Fig. 3).

**Figure 1:**
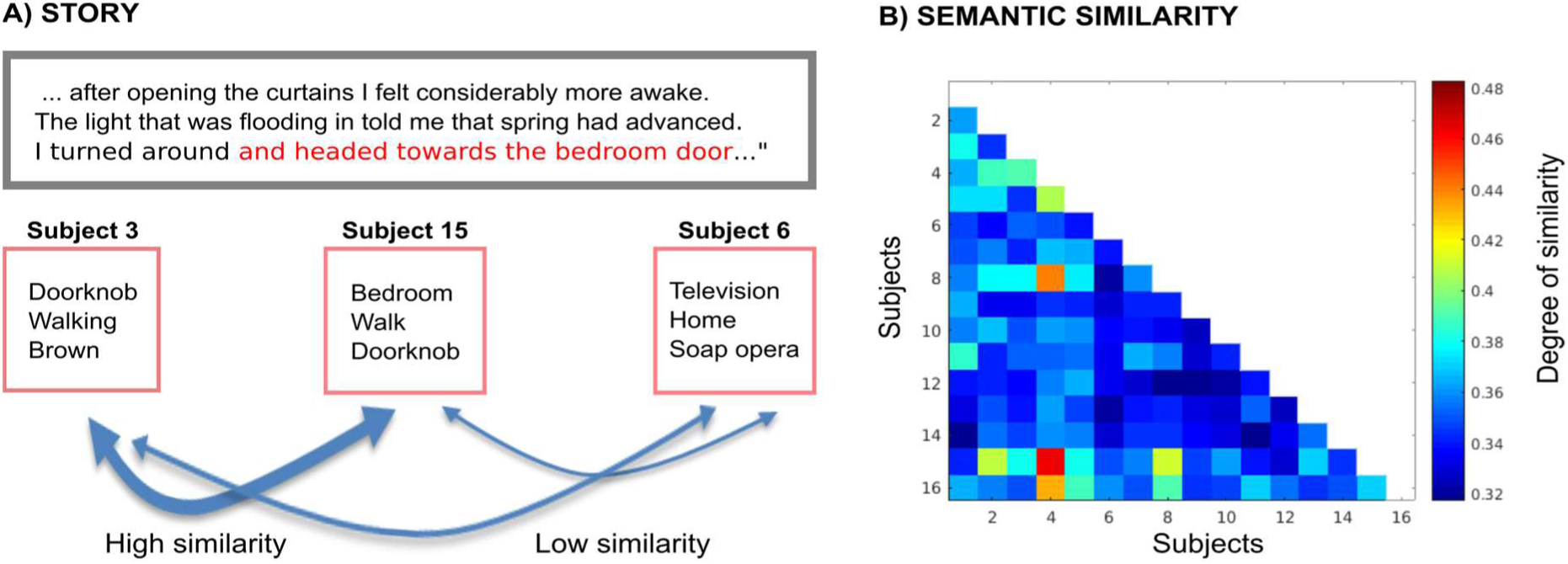
Similarities/differences of subjects’ individual semantics when listening to the narrative. LEFT: Excerpt from the narrative with one phrase-segment highlighted with red font color. Word lists produced by three representative subjects to this particular segment are shown below as examples of similarities and differences in the individual semantics (note that both the narrative excerpt and word lists have been here translated to English for illustration purposes). RIGHT: Correlation matrix showing LSA- and WordNet-derived similarities/differences of subjects’ individual semantics when listening the narrative. While some subject pairs exhibit striking similarity, there were also robust differences across many subject pairs. Note that the values plotted here mark mean subject pair-wise similarities-differences across the whole narrative.

**Figure 2:**
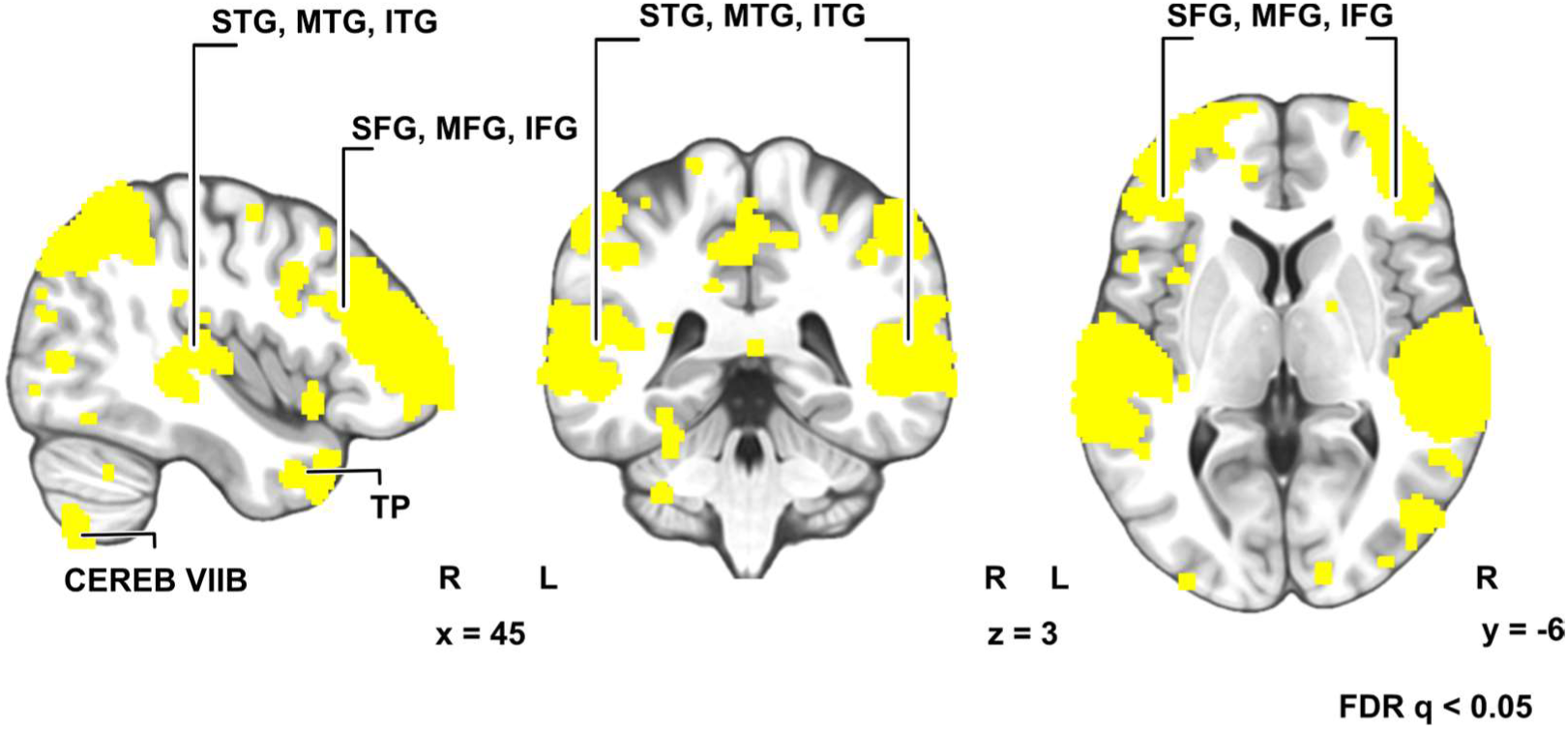
Inter-subject correlation (ISC) of BOLD signals (FRD corrected q< 0.05).

**Figure 3:**
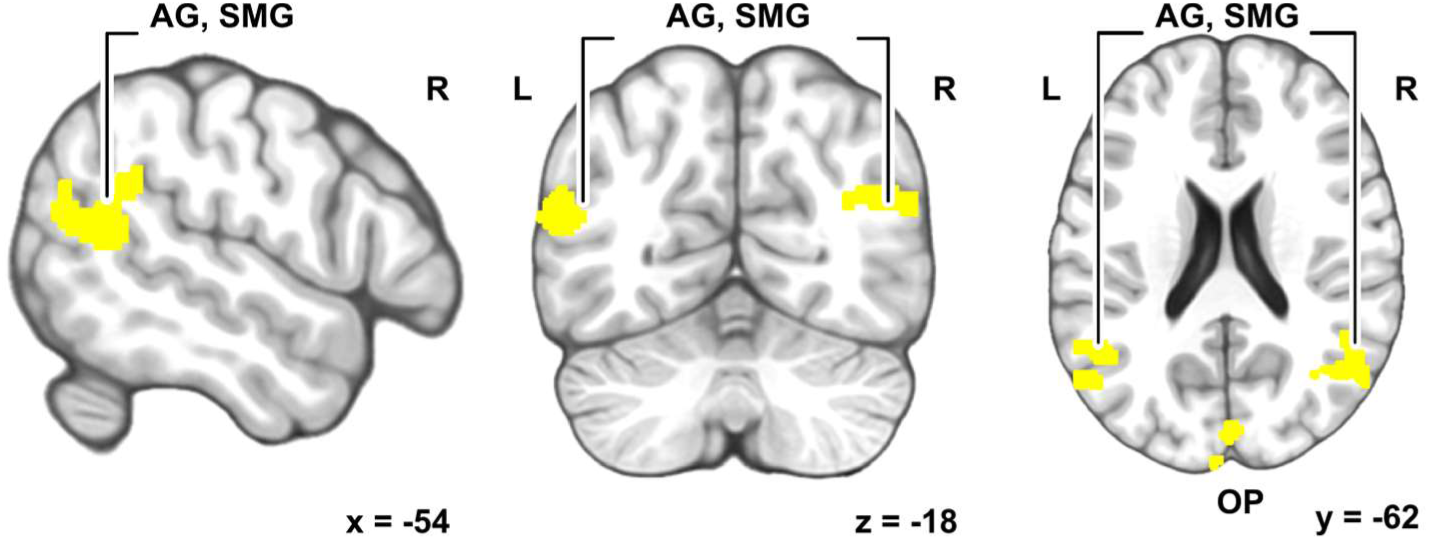
Brain areas where similarities in perceived semantics of the narrative significantly predicted inter-subject similarity of brain activity during narrative listening. (AG = angular gyrus; SMG = supramarginal gyrus; OP = occipital pole). Peak activation (see supplementary info) at left supramarginal gyrus −56, −50, 26, right angular gyrus 48, −62, 26, and right cuneus 4, −88, 18. Unthresholded correlation-value maps from the RSA analysis can be found in 3-D brain space at Neurovault (https://neurovault.org/collections/KCKVHDCV/).

## Discussion

When listening to a captivating story we often can virtually see the beautiful scenes, various objects, and protagonists acting in their environment^19^. Such immersion in the flow of a story is a unique human ability made possible by the brain seamlessly calling upon one’s own past experiences and acquired generic knowledge to give rise to the vivid mental contents in the form of associations^11^ and associated mental imagery^6^. In the present study, the word lists produced by the subjects describing what had come to their minds while listening to the narrative showed robust between-subject variability, suggesting that the triggered mental contents are individualistic. While previous studies have shown inter-individual differences in, e.g., associations elicited during viewing of pictures^11^, we present here, to our knowledge novel, methodology to measure and analyze differences in semantics and associated mental imagery elicited by a narrative.

Listening to the narrative elicited significant ISC in extensive set of brain bilaterally (**Fig. 2**). Similarity of activation extended beyond the classical linguistic areas to bilateral frontal, temporoparietal brain areas, extending to midline regions such as precuneus and cuneus and right cerebellum. These results are highly similar to findings in previous studies using naturalistic linguistic stimuli^20–22^, however, significant ISC does not *per se* reveal brain regions supporting semantics and associated mental imagery elicited by the narrative as significant ISC can be due to similarity in a variety of other cognitive and processes that take place during narrative listening besides semantic processing and imagery.

Notably, between-subject similarities in perceived semantics of the story predicted between-subject similarities in local brain hemodynamic activity in inferior parietal lobule (SMG and AG) as well as in cuneus in visual cortex. SMG and AG belong to the semantic network laid out in a previous meta-analysis of the semantic system of human brain^1,2^ and, supporting recent observations about semantic representations in both left and right hemispheres^3^, we observed the similarity of activity bilaterally. It has been suggested that areas in the inferior parietal lobe function as convergence zones for concepts and event knowledge, and they receive input from sensory, action, and emotion systems^2^. However, SMG is also activated by complex motor sequences such as articulation^23^, and phonological processing^24^, and it’s activity has been identified in conditions that pose specific challenge for semantic prosessing^25^. Instead, AG has been shown to be involved in both semantic processing^1,25^ and autobiographical memory, which, in fact, has been suggested to build on general semantic memory processing. Importantly, AG has been found to serve as a hub in integrating semantic information into coherent representations^26,27^, and structural differences in the area have been found to be related to inter-individual differences in a task that requires combining concepts^28^. Moreover, given that the heteromodal AG has been indicated to take part in a variety of cognitive functions^29,30^, the involvement of AG in building individualized semantics and integrating visual processes is plausible.

Similarity of associations predicted similarity of brain activity also in early visual areas (**Fig. 3**), a finding that is in line with previous research suggesting that visual imagery is supported by same areas as visual perception. Results of the current study, therefore, suggest that the narrative may have elicited similar mental imagery for individuals using semantically more similar words to describe what came to their minds during listening of the narrative^4^. This would not, of course, necessarily imply identical mental images, but rather similarity in the process in which the individuals engaged in generation of the mental imagery during listening to a story. Thus, one can speculate whether individuals with more similar activity in early visual areas drew upon visual information stored in the brain related to objects, scenes, and events in the narrative in similar accuracy or strength^8^.

The practical limitation of our method is that it is highly laborious for experimental subjects to report associations once every 3–5 s for narratives longer than the eight minute one used in the present study. Given this, it is also possible that we might have been able to observe significant activity in some other areas of the semantic network in the present study had we been able to collect more data. Thus, while it can be safely concluded that the inferior parietal and visual cortical areas are involved in generation of individualized semantics and associated mental imagery, one should exercise caution against concluding that some other areas would not be involved in this process. Nonetheless, while ISC quantifies the similarities across individuals, our approach constitutes a methodological advance that makes it possible to quantify inter-individual differences of semantic representations and mental imagery during narrative listening in the human brain.

In conclusion, individuals with more similar activity SMG and AG of the inferior parietal lobe, as well as in early visual cortical areas, specifically cuneus, during listening to a narrative also elicited mental associations that were semantically more similar. Thus, these areas seem to support elicitation of individual meanings and flow of mental imagery during listening to a captivating narrative.

## Materials and Methods

The current study reports data from 16 healthy female volunteers (ages 20 - 42, all right-handed according to Edinburgh handedness inventory^31^) that first listened or read the narrative. The behavioral and fMRI data for the current experiment was obtained in parallel with a broader-scope fMRI experiment (N=29) investigating brain mechanisms during listening, reading, and lipreading a narrative in a counter-balanced order. Only subjects who listened or read the narrative first were included in the present study and asked to produce the associations. The narrative described, from first-person perspective, daily events in a life of a woman (for original Finnish and English-translated versions of the story, see Appendix A and B **below**). Furthermore, a written, 614-word transcript of the narrative was created. During fMRI scanning, written words were presented simultaneously centrally on the screen time-locked to each word of the original spoken narrative. In those cases where the duration of the words in the spoken narrative were very short, two or three words were presented simultaneously to keep the timing.

Presentation software (Neurobehavioral Systems Inc., Albany, California, USA) was used for presenting the stimuli. The audio stimuli were played with an MRI-compatible in-ear earbuds (Sensimetrics S14 insert earphones). In addition, MRI-safe protecting earmuffs were placed over the earbuds for noise removal and safety. Before the actual experiment, sound intensity was adjusted for each subject to be loud enough to be heard over the scanner noise by playing example stimuli that were normalized to the same level as the auditory stories during a dummy EPI sequence before. In the MRI scanner, the stimulus videos and texts were back-projected on a semitransparent screen, using a Panasonic PT-DZ110XEJ projector (Panasonic Corporation, Osaka, Japan). The viewing distance was 35 cm.

During narrative presentation the subjects’ brain hemodynamic activity was recorded with fMRI (Siemens 3-Tesla Skyra, Erlangen, Germany; standard 20-channel receiving head/neck coil; T2-weighted echo-planar imaging sequence with 1700 ms repetition time, 24 ms echo time, flip angle 70°, each volume 33 × 4 mm slices, martix size 202 × 202 mm, in plane resolution 3 × 3 mm) at the Advanced Magnetic Imaging Centre of the Aalto University. Anatomical T1-weighted structural images were acquired with 1 × 1 × 1 mm resolution (MPRAGE pulse sequence, TR 2530 ms, TE 3.3 ms, TI 1100 ms, flip angle 7°, 256 × 256 matrix, 176 sagittal slices). These fMRI data were analyzed by calculating voxel-wise inter-subject correlations (ISC) of hemodynamic activity using our in-house BraMiLa analysis pipeline based on a published toolbox^17^. After the fMRI session, the subjects were presented the narrative again in writing, divided to 128 consecutive coherent phrases (3-5 sec in duration), and were instructed to write words (within 20–30 sec) best describing what came to their mind upon hearing each segment in fMRI. After removing the conjunctions from the words, and translating words into English, we then utilized latent semantic analysis^12^ (LSA; implemented using Gensim Python library^32^) combined with the WordNet^13^ knowledge on the content words. LSA assumes that words that occur in the same context have similar meanings. We used European Parliamentary corpus database^33^ to produce a word co-occurrence statistic which was turned into a 300-dimensional^34^ semantic space through singular value decomposition (SVD). Each word list produced by the subjects was represented as a vector in this semantic space and the similarity between word lists was computed as the cosine similarity of the vectors. This LSA-derived similarity was increased using WordNet knowledge e.g. when the words are in the same WordNet synset or one word is the direct hypernym of the other^18^.

The subject pair-wise similarities in semantics were then utilized to predict similarities in brain function for each voxel to derive statistical parametric maps that were corrected for multiple comparisons (cluster forming threshold p=0.05, cluster extent threshold 125 voxels). The behavioral and fMRI data for the current experiment was obtained in parallel with a broader-scope fMRI experiment (N = 29) investigating brain mechanisms during listening, reading, and lipreading a narrative, as well as an unintelligible, gibberish version of the same narrative. The stimulus sequence in the full experimental design consisted of six different narratives, three intact (lipread, read and listened), and three gibberish versions of the same narratives. The word-list associations were obtained only from a subset of subjects reported here. The lipreading results will be reported separately^35^.

## Acknowledgments

We kindly thank the subjects for their participation in the experiment. We also thank personnel of imaging facilities in AMI centre of Aalto University School of Science, Espoo Finland, and especially nurse Marita Kattelus for her help in fMRI imaging.

## Author contributions

SS planned the experiment, prepared stimuli, collected and analyzed data, prepared figures, wrote the manuscript as first author.

JA planned the experiment, collected and analyzed data, prepared figures, wrote the manuscript. EG analyzed data, prepared figures, wrote the manuscript.

MK analyzed data

TH planned the experiment and supervised data analysis

MS planned the experiment, supervised collecting data, wrote the manuscript

MB planned the experiment, supervised data analysis, wrote the manuscript

IPJ planned the experiment, supervised data analysis, prepared figures, wrote the manuscript

## Data and materials availability

Unthresholded correlation-value maps from the RSA analysis can be found in 3-D brain space at Neurovault (/collections/KCKVHDCV/)

